# Thermodynamic Analysis of Protein-Nanoparticle Interactions Links Binding Affinity and Structural Stability

**DOI:** 10.1101/2025.08.21.671591

**Authors:** Chathuri S. Kariyawasam, Radha P. Somarathne, Naomi C. Hellard, Nicholas C. Fitzkee

**Author notes:** To whom correspondence should be addressed: Dr. Nicholas C. Fitzkee Department of Chemistry, Mississippi State University 310 President Circle Mississippi State, MS 39762.

## Abstract

When nanoparticles and nanoplastics enter biological fluids, their surfaces are rapidly coated with proteins, forming a corona that governs biological responses. However, understanding protein-surface interaction energetics remains a significant challenge. Here, we examine how protein charge distribution affects adsorption to polystyrene nanoparticles (PSNPs) by generating a series of lysine-to-alanine variants of the GB3 protein. Using isothermal titration calorimetry (ITC), we found that the K19A variant binds most strongly to both non-functionalized and carboxylate-functionalized PSNPs. ITC thermograms indicate that K19A forms a stable monolayer, while other variants exhibit multilayer adsorption. We hypothesize that removing lysine at position 19 creates a flatter, more neutral interaction surface that promotes efficient initial binding. Fluorescence denaturation experiments show that PSNPs destabilize GB3 protein variants, and binding correlates strongly with protein unfolding (r = 0.82, p < 0.01 for COOH-PSNPs and r = 0.76, p < 0.03 for non-functionalized PSNPs). These results reveal how protein stability and charge distribution shape adsorption thermodynamics, offering a framework for predicting protein-surface interactions.

**TOC Image:** 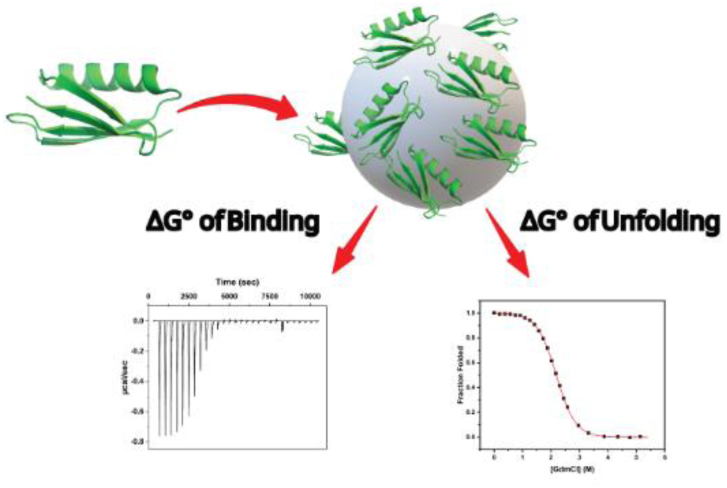

## Introduction

The interaction between proteins and polymeric surfaces is a critical area of study, with far-reaching implications for understanding how organisms respond to nano- and microplastics. When these materials enter biological environments, they are rapidly coated with biomolecules, forming a complex known as the protein corona.^1–3^ This corona significantly influences the particle’s biological identity and subsequent interactions within the body.^4^

Rather than being a uniform layer, the corona is a complex, dynamic structure composed ot two components: the hard corona and the soft corona. The hard corona consists of proteins that bind tightly to the surface with high affinity and long residence times, often undergoing conformational changes, making them difficult to remove. The soft corona contains proteins that associate loosely and exchange rapidly with the surrounding medium.^5–8^ The formation of these layers is time-dependent. Initially, abundant proteins with high association rates adsorb to the surface, but less abundant, higher-affinity proteins can later displace them. This process is known as the Vroman effect.^9^ As a result, the hard corona typically stabilizes over time, while the soft corona remains in dynamic equilibrium.^3,10^

Understanding the formation and composition of both corona components is crucial because they collectively govern how nano- and microplastics interact with cells, tissues, and organs. Studies of polystyrene microplastics have been challenging because of their poor colloidal stability,^11^ but the coronas formed on stable polystyrene nanoparticles (PSNPs) have been studied extensively. The hard corona often determines the particle’s long-term biological identity,^6^ while the soft corona can dominate early interactions and modulate the evolution of the hard corona over time.^6,12,13^ Additionally, the corona affects how the immune system recognizes and responds to nanoparticles: certain proteins promote immune recognition, while others facilitate immune evasion.^12^ The contrasting stabilities of the hard and soft coronas therefore shape the biological responses over different timescales.^14,15^

A key open question is how the energetics of surface binding relate to a protein’s structure and stability. Prior work has provided valuable but sometimes inconsistent insights. Cedervall *et al.*used HSA and fibrinogen to demonstrate that nanoparticle surface properties like hydrophobicity and curvature influence binding kinetics, residence times, and conformational changes.^16^ Milani *et al.*, using fluorescence correlation spectroscopy of transferrin, revealed two distinct adsorption timescales, corresponding to hard and soft corona formation. Their work quantified the transition from monolayer to multilayer coverage and demonstrated the importance of the protein-to-nanoparticle ratio.^8^ Kelly *et al.* emphasized the importance of biomolecular recognition and specific peptide motifs in defining protein orientation and downstream biological outcomes like intracellular trafficking.^17^ Together, these studies suggest that protein-surface interactions involve heterogeneous affinities, but they differ in how they interpret the source of this heterogeneity: Cedervall *et al.* emphasize nanoparticle surface properties, Milani *et al*. highlight protein exchange dynamics, and Kelly *et al.* focus on recognition patterns. Moreover, they propose different implications for how proteins orient on the nanoparticle surface.

One challenge in reconciling these and other perspectives is the diversity of proteins studied. Albumin, transferrin, and fibrinogen are important serum proteins, but vary widely in shape, size, and stability. This variation complicates systematic interpretation. A promising alternative is to complement serum protein studies with a mutagenesis strategy that examines the same model protein while systematically altering a small number of residues.

To address this gap, we combine site-directed mutagenesis with biophysical measurements to probe how protein stability and charge distribution influence surface binding. Mutagenesis is a powerful tool for dissecting amino acid contribution to protein behavior,^18–20^ but it has rarely been applied to protein-nanoparticle systems. In this study, we use the third IgG binding domain of the streptococcal protein G (GB3) as a model system,^21^ introducing lysine-to-alanine mutations to investigate how electrostatic interactions affect both binding affinity to PSNPs and folding stability in their presence. GB3 is an ideal model: its small size (56 amino acids), well-characterized structure, and high lysine content (seven residues) make it particularly sensitive to charge redistribution.^22–26^ As an extracellular domain, GB3 can also mediate bacterial interactions with plastic particles in the environment.^26,27^

In addition, we examine how nanoparticle surface chemistry – carboxylate-functionalized (COOH) vs. non-functionalized (NF) PSNPs – modulates binding thermodynamics and structural effects. Our results show that variants with the strongest binding affinities are also the most destabilized upon adsorption. This correlation suggests that proteins most tightly bound to polystyrene surfaces are also the most structurally perturbed, a finding that complicates predictions of protein structure in nanoparticle coronas and potentially other surface adsorption phenomena.

## Materials and Methods

### Protein Expression and Purification

All the GB3 lysine variants were expressed and purified as described previously.^25^ Briefly, the BL21 cells were grown in LB medium overnight at 37 °C and transferred into 1L of ^15^N M9 media to reach an initial OD_600_ of 0.05. The cells were incubated at 37 °C at 200 rpm and induced with a final concentration of 0.5 mM isopropyl β-D-1-thiogalactopyranoside (IPTG) when the cells reached an OD_600_ of 0.6. Then, the cells were allowed to grow for an additional 6 hours. The cells were harvested by centrifugation at 6,500 rcf for 30 min. The cell pellet was then resuspended in 20 mL of cold lysis buffer (20 mM NaH_2_PO_4_ pH 7.5, 50 mM NaCl, 5 mM EDTA, 0.5 mg/mL lysozyme) and rocked on ice for 30 min. The resuspended cell pellet was then sonicated on ice in a FisherBrand sonicator at 45% power for 6 minutes (30 s pulse on, 30 s pulse off). Next, the lysed cells were incubated in an 85 °C water bath for 15 min, swirling every 3-4 minutes. The solution was cooled rapidly in an ice water bath for 30 min, and streptomycin sulfate was added to a final concentration of 1.0% (w/v) to precipitate DNA. This solution was allowed to sit for an additional 10 min on ice. The cell debris, precipitated proteins, and DNA were removed by centrifugation at 80,000 rcf for 45 min. The supernatant was collected and purified in an AKTA FPLC with a HiTrap Q FF anion exchange column. In this step, 50 mM NaCl, 20 mM NaH_2_PO_4_ pH 4.5 was used as the wash buffer and 1 M NaCl 20 mM NaH_2_PO_4_ pH 4.5 was used as the elution buffer. Under these conditions, the impurities stick to the column and the GB3 comes in the flow-through. This fraction was concentrated with 3.0 kDa EMD Millipore Amicon Ultra centrifugal filters (Pall Corporation) and loaded to a HiLoad 26/600 Superdex 75 pg gel filtration column equilibrated with gel filtration buffer (50 mM NaCl, 20 mM NaH_2_PO_4_ pH 6.5). GB3 eluted at around 220 mL, and the fractions were analyzed by gel electrophoresis for purity.

### Characterization of Nanoparticles

Non-functionalized polystyrene nanoparticles (NF-PSNPs) and carboxylate-functionalized polystyrene nanoparticles (COOH-PSNPs) with a diameter of 50 nm were purchased from PolySciences, Inc. (catalog numbers 08691 and 15913, respectively). The hydrodynamic diameter and the zeta potential of the nanoparticles were measured using an Anton Paar Litesizer 500 DLS at 25 °C. The concentration of the nanoparticle stock solution was calculated using the manufacturer’s particle count and verified using gravimetric analysis. All the nanoparticles were extensively dialyzed against water and then the buffered solution (20 mM NaH_2_PO_4_, 50 mM NaCl, pH 6.5) to remove surfactants before use in experiments. Removal of surfactants was confirmed using NMR and pendant drop analysis techniques.^28,29^ The surface tension of nanoparticles before dialyzing (54 mN/m) and after dialyzing in water (70 mN/m) was measured and compared with that of pure water (72 mN/m).

### Isothermal Titration Calorimetry

Isothermal titration calorimetry (ITC) measurements were performed on a Microcal VP-ITC instrument (GE Healthcare). All protein and nanoparticle solutions were dialyzed in 20 mM H_2_NaPO_4_, 50 mM NaCl, pH 6.5 buffer (NP binding buffer). A typical ITC titration experiment with COOH-PSNPs involved 35 injections of a particular GB3 variant as the titrant (8 μL per injection from a 500 μM stock) at 360 s intervals into the sample cell (volume = 1.4 mL) containing 175 nM COOH-PSNP solution. A typical titration experiment with NF-PSNPs involved 28 injections of each GB3 variant (10 μL per injection from a 200 μM stock) at 360 s intervals into the sample cell containing the 25 nM NF-PSNP solution. All the experiments were performed at 25 °C, and the sample cell was continuously stirred at 480 rpm during the experiments. Control experiments were performed to determine the heats of dilution and subtracted from the initial titration data for each experiment. Baseline corrections for the raw thermograms were performed using NITPIC software,^30^ and the data were analyzed using the CHASM software^31^ to obtain the binding stoichiometry (*N*), binding affinity (*K*_a_), and other thermodynamic parameters by fitting with two sets of identical sites model. All the reported parameters are an average of three independently run experiments, and error bars are calculated as the standard error of the mean (SEM).

### Fluorescence Denaturation Experiments

Fluorescence measurements were carried out on a Horiba Fluormax 4 fluorescence spectrophotometer using a 1.00 cm path-length quartz cell. Samples were prepared with 4 μM GB3 variants and 1.5 nM NF-PSNPs and 2.5 μM GB3 variants and 1.5 nM COOH-PSNPs in NP binding buffer. Guanidinium chloride (GdmCl) was used as the denaturant. Optimized injections of GdmCl with a nominal stock concentration of 7.7 M were titrated into all samples with a 6 min interval between injections for equilibration. Actual GdmCl concentrations were determined using refractometry.^32^ A constant sample volume was maintained throughout the experiment by using an auto-titrator (Hamilton Microlab 600) interfaced with the Fluoromax software. The sample was stirred at 350 rpm throughout the experiment. The emission spectra were recorded in the wavelength range of 305-400 nm upon excitation at 295 nm using a 3 nm slit width. Appropriate blank experiments were conducted and subtracted from the spectra before analyzing the data. The free energies of unfolding (Δ_unfold_G°) for proteins in the absence and presence of PSNPs were calculated by fitting the linear extrapolation model.^33^ Fitting scripts are available at https://github.com/FitzkeeLab/fluor_denaturation.

### Circular Dichroism Experiments

The circular dichroism (CD) measurements were performed on a Jasco 1500 CD spectropolarimeter at 25 °C using a 0.2 mm pathlength demountable quartz cuvette. The CD spectra were recorded in the wavelength range of 185-260 nm with a scan rate of 20 nm/min and 16 sec integration time. The concentration of GB3 variants was 40 μM and 10 nM NF-PSNPs was used in NP binding buffer (see ITC section).

### T_1ρ_ relaxation Experiments

^15^N HSQC-resolved T_1ρ_ relaxation experiments were carried out at 25 °C on a 600 MHz Bruker Avance III cryoprobe-equipped NMR spectrometer.^34^ The ^15^N spin lock field was 2 kHz. The relaxation experiments for the protein in the presence and absence of NPs were carried out with 8 different delays (2, 20, 50, 100, 160, 190, 210, and 290 ms). Samples were prepared using 150 µM GB3 variants and 120 nM PSNPs in a buffer solution containing nanoparticle binding buffer and 6% D_2_O. The T_1ρ_ values were not converted to T_2_ values but were compared (with and without nanoparticles) directly.

## Results

### Systematic Mutagenesis Reveals Differences in PSNP Binding of GB3 Variants

Previously, our work has demonstrated that lysine residues play an important role in GB3 binding to gold nanoparticles (AuNPs, **Figure 1A**).^23,25^ This observation led us to investigate GB3 lysine to alanine variants and their interaction with polystyrene surfaces. Changing lysine to alanine alters the electrostatic surface of GB3 (**Figure 1B**), and we generated all possible variants for this study (K4A, K10A, K13A, K19A, K28A, K31A, and K50A). Two types of polystyrene nanoparticles (PSNPs) were used (nominal diameter 50 nm) to explore the effects of surface interactions. Both COOH-PSNPs and NF-PSNPs possessed a negative zeta potential. While this observation is expected for COOH-functionalized PSNPs, non-functionalized PSNPs exhibit a negative zeta potential due to sulfonate groups introduced during PSNP synthesis (**Table S1, Supporting Information**).^5^ Initially, we hypothesized that the GB3-PSNP interaction would have a similar binding pattern to what was observed for citrate-capped AuNPs **(Figure 1C**). When bound to AuNPs, GB3 exhibits essentially no line broadening in NMR titrations, reflecting stable, kinetically controlled adsorption.^22,24,25^ In contrast, GB3 binding to PSNPs exhibits significant line broadening (Δ*R*_2_), reflecting a dynamic exchange between free and surface-bound protein (**Figure 2A**).^35,36^ This is consistent with other studies of PSNP-protein interactions, where the dynamic exchange reflects a soft corona formation.^1,8,37,38^ Dynamic light scattering (DLS) experiments on mixtures of GB3 variants and PSNPs also revealed an increase in apparent hydrodynamic diameter, indicating the formation of a protein corona (**Figure 2B**); however, no clear trend in binding was observed. Together, these results led us to explore isothermal titration calorimetry (ITC)^39^ as a technique that could potentially differentiate GB3 binding to PSNPs over the set of lysine to alanine variants.

**Figure 1.**
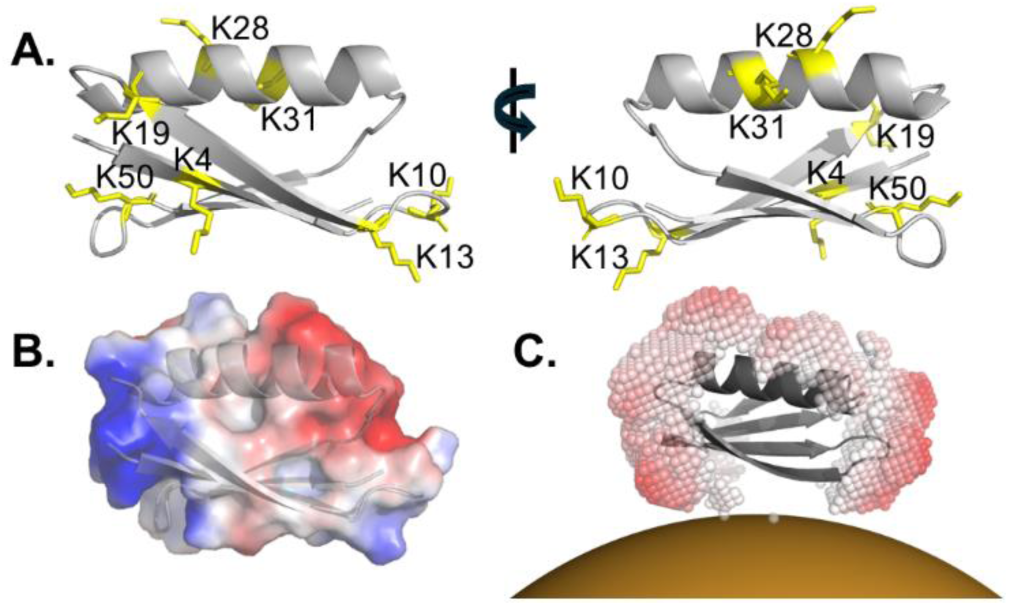
Illustration of the (A) Lysine residues and (B) Electrostatic surface of GB3 as calculated by APBS^40^, (C) GB3 binding to AuNPs representing the binding surface using color-coded virtual atoms, colored white (low affinity) to red (high affinity) using the surface-binding algorithm calculated by Xu *et al.*^25^

**Figure 2.**
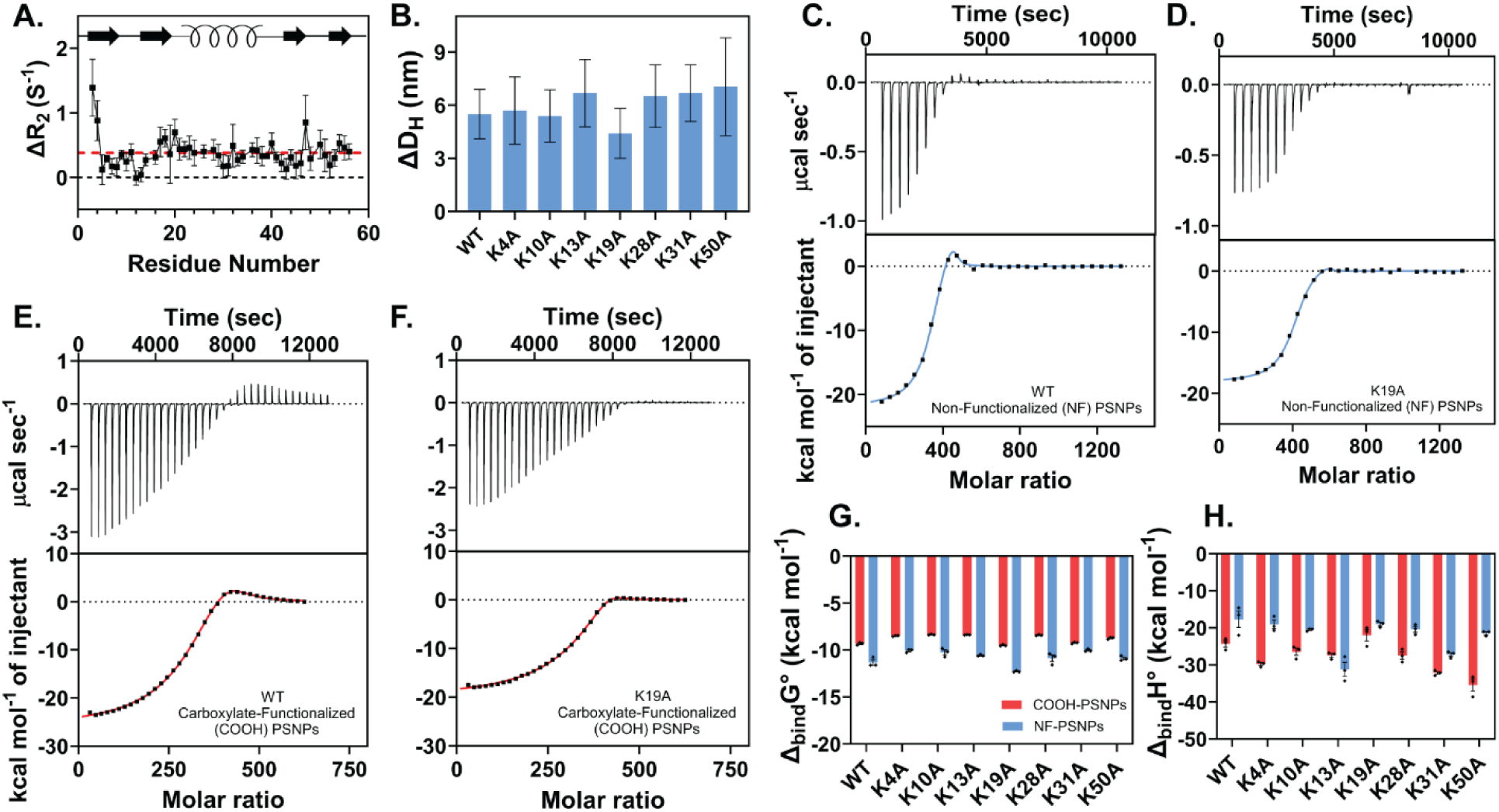
(A) The Δ*R*_2_ values for WT GB3 measured with and without PSNPs using T_1ρ_ relaxation NMR experiments (B) The change in hydrodynamic diameter of PSNPs due to formation of a protein corona. (C-F) Representative ITC data with NF-PSNPs for (C) WT GB3 and (D) K19A GB3 and with COOH-PSNPs for (E) WT GB3 and (F) K19A GB3. (G, H) ITC parameters are plotted for the free energy of binding (Δ_bind_G°, G) and enthalpy of binding (Δ_bind_H°, H) for COOH- PSNPs (red) and NF-PSNPs (blue). Error bars represent the standard error of the mean (SEM) from three independent measurements.

Using ITC to determine the binding parameters of these lysine variants revealed significant differences, both among the variants themselves and between COOH- and NF-PSNPs (**Figure 2**). The thermograms of all variants featured a similar behavior, except for K19A. (**Figure 2C, D, E, F**) The interaction of K19A was consistently exothermic until saturation, while the interaction of other variants was exothermic at the beginning and gradually changed to endothermic as saturation was approached, resulting in a hump-like feature in their isotherms. This feature indicates that the GB3 binding to PSNPs consists of at least two distinct thermodynamic signatures. We extracted the thermodynamic parameters using a model with two sets of N identical sites (**Table S2 and S3, Supporting Information**). Two sets of sites were also needed for fitting the K19A data, even though a distinct endothermic feature was not seen for this variant. We refer to the strong, exothermic binding as Process 1; and the endothermic process is referred to as Process 2. Similar features were observed in ITC experiments performed by Viola *et al.* when monitoring the interaction of tau protein with ultra-small AuNPs. They suggest that this feature might be due to the multilayer binding of proteins to NPs.^41^ In their explanation, the first binding process is dominated by the exothermic binding of the protein to the ultra-small AuNP surface. The second binding process occurs after the initial layer of protein is formed; in this case, the endothermic process results from hydrophobic interactions on the particle surface, as hydrophobic interactions are commonly entropically driven at room temperature.^42^ For all variants measured, Process 1 is always stronger than Process 2 and typically involves more proteins, which is consistent with an initial layer of formation followed by weaker multiple-layer binding.

To facilitate comparative analysis across variants, we focused on thermodynamic parameters from Process 1, the initial, exothermic binding process. Analyzing the ITC data, we observed that the K19A variant showed the highest free energy of binding (Δ_bind_G°) to both types of NPs, indicating the strongest affinity. The WT protein showed the next highest binding affinity for both NF-PSNPs and COOH-PSNPs. Among the other variants, the K31A variant displayed a higher affinity for COOH-PSNPs but a lower affinity for NF-PSNPs compared to the other variants. Conversely, the K28A variant demonstrated a higher affinity for NF-PSNPs and a lower affinity for COOH-PSNPs relative to the other variants. Notably, all the lysine-to-alanine variants exhibited higher binding affinities for the NF-PSNPs compared to the COOH-PSNPs (**Figure 2G**). At the pH where experiments were performed (pH 6.5), all GB3 variants are expected to possess a net negative charge, making binding to NF-PSNPs slightly more favorable.

In contrast, when examining the binding enthalpies, they were generally more negative for the COOH-PSNPs compared to the NF-PSNPs. The one exception to this trend is the K13A variant, which had a slightly more favorable enthalpy for NF-PSNPs than COOH-PSNPs. The K19A and WT variants, which had the highest binding affinities, exhibited the least negative binding enthalpies for both NP types. However, the trends observed in the free energies of binding for the other variants did not directly follow the trends seen in the binding enthalpies (**Figure 2H**). Overall, the calorimetric profiles of protein variants suggest that GB3-PSNP binding has multiple thermodynamic signatures, each with its own characteristic binding interactions.

### Evidence for Multilayer Binding to PSNPs

Intrigued by the behavior of the K19A variant, we sought to validate whether the observed hump in the ITC data indeed originates from multilayer protein binding to NPs. We focused our investigation on the protein corona formed on COOH-PSNPs, as these particles exhibited a more pronounced endothermic feature in their thermograms. We performed dialysis on PSNP samples that had already been saturated with protein. This dialysis reduces the concentration of free GB3 in solution, shifting the equilibrium so that loosely bound proteins desorb from the NP. A 12 kDa molecular weight cutoff (MWCO) was used, which dilutes GB3 while retaining NPs in the dialysis bag. These samples were then titrated again in the ITC with GB3 (**Figure 3A**). For all samples except K19A, the endothermic hump was observed with very little (or no) exothermic heat (**Figure 3B and Figure S1, Supporting Information**). This suggests that, during titration, most of the (exothermic) NP surface sites are occupied; these fill up early in the titration, and additional protein added contributes to the second, endothermic thermodynamic feature. This supports the idea that the endothermic hump results from late binding to the PSNP, as would be expected if multiple layers were forming. Notably, no endothermic features are observed for K19A, suggesting the absence of multilayer binding (**Figure 3C**). Overall, the integrated endothermic enthalpies were similar for the dialyzed samples and the complete ITC titrations (compare **Figure 2E** vs. **Figure 3B**); however, direct comparison of these values is challenging for several reasons. First, after dialysis, the starting protein concentration and degree of PSNP saturation is unclear. Second, the PSNP concentration itself is difficult to know without destructive analytical techniques. Thus, the thermograms after dialysis were not fit with a binding model, and instead a qualitative interpretation was used.

**Figure 3.**
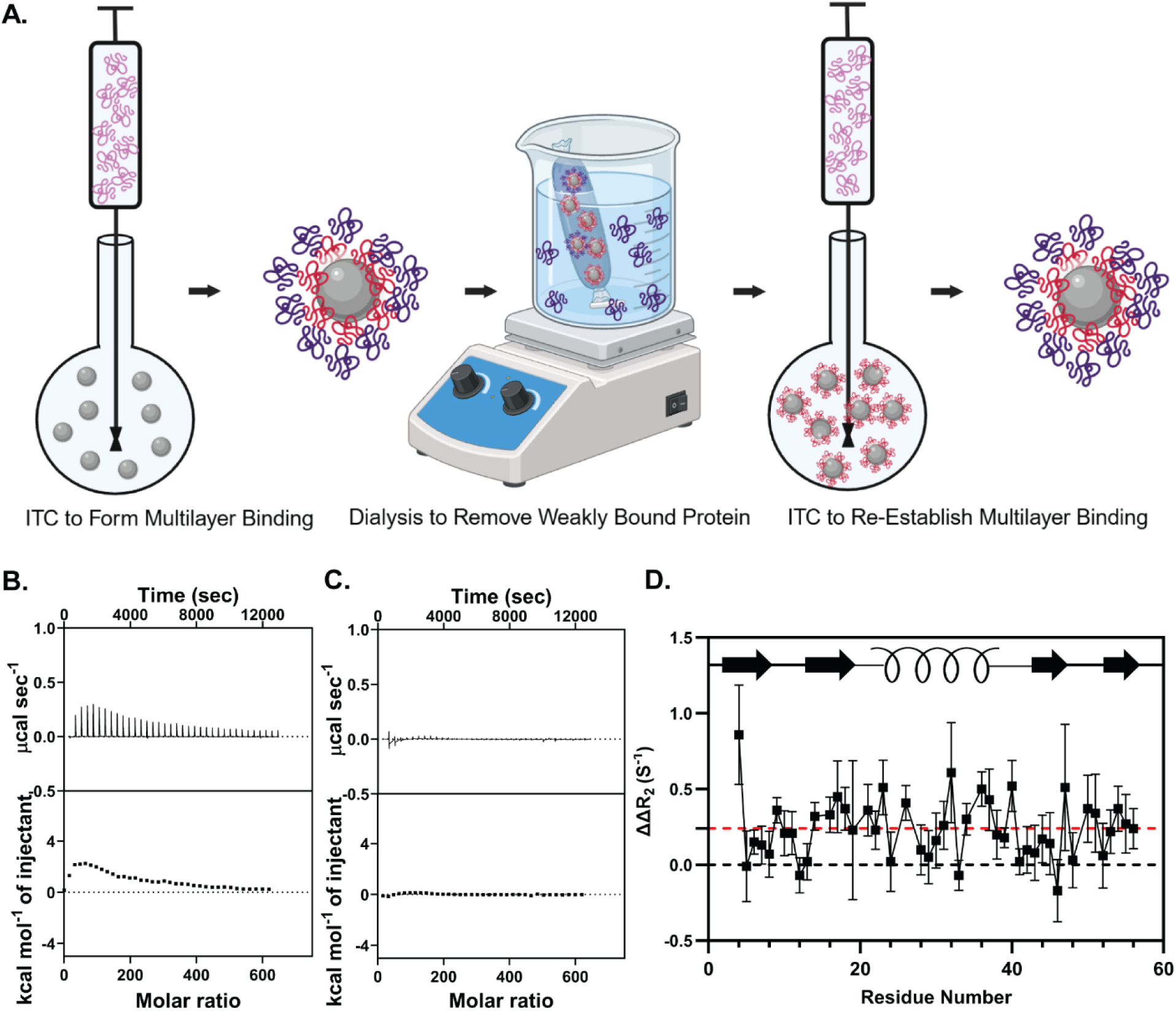
Evidence for multilayer binding in non-K19A GB3 variants. (A) A schematic representation of the steps involved in re-establishing multilayer protein binding in the ITC. (B, C) Representative ITC thermograms of binding after removal of weakly bound proteins for (B) WT GB3 and (C) K19A GB3. (D) The difference in ΔR_2_ rates of WT GB3 and K19A GB3. This difference indicates a positive ΔΔR_2_ due to enhanced lifetime line broadening, reflecting multilayer exchange in WT GB3 bound to NF-PSNPs.

We sought to use DLS as a complementary technique to visualize multilayer binding in protein variants. While suggestive differences were observed between K19A and the other variants (**Figure 2B**), there was no statistically significant difference detected (p = 0.24). This is likely because DLS is not sensitive to subtle changes in dynamically exchanging protein coronas.^43,44^ Instead, we used an NMR-based approach to monitor adsorption and desorption among the variants. In the presence of NPs, dynamic binding contributes to lifetime line broadening, an additional exchange contribution to transverse relaxation rates.^45,46^ This effect manifests itself as an additional contribution to the R_2_ rate, or small, positive values of Δ*R*_2_for relaxation rates measured with and without nanoparticles. The Δ*R*_2_ values were compared for WT and K19A variants in the presence and absence of NF-PSNPs. The Δ*R*_2_ values for WT GB3 were consistently higher than those for K19A GB3, which is consistent with increased surface exchange in WT GB3 (p < 2.2 × 10^−10^) indicating a positive ΔΔ*R*_2_ rate (**Figure 3D, Figure S2 and S3, Supporting Information**). Thus, while K19A exhibits Δ*R*_2_ values consistent with binding PSNPs, other GB3 lysine variants experience enhanced exchange, which supports a model of multilayer binding. Together, the dialysis experiments and enhanced lifetime line broadening support the hypothesis that the endothermic hump in ITC experiments corresponds to weaker, multilayer binding, providing an explanation for the thermodynamic parameters of Process 2 (**Tables S2 and S3, Supporting Information**).

### Structural Perturbations of Bound GB3 Variants

The strong binding of Process 1 is characterized by a broad range of enthalpies, even though the values of Δ_bind_G° are fairly consistent. For example, consider that the range of Δ_bind_G° values on NF-PSNPs spans 2-3 kcal mol^-1^, whereas the range of Δ_bind_H° spans nearly 20 kcal mol^-1^ (**Figure 2G-H**). The behavior is characteristic of enthalpy-entropy compensation, which is often associated with the hydrophobic effect.^47^ Therefore, we sought to investigate whether structural changes (such as protein unfolding) were responsible for differences between lysine-to-alanine variants. As an initial step, we sought to compare whether the folded protein structure could predict the enthalpy or free energy of binding. We used a model that we developed previously, where basic residues on the folded protein’s surface were used to identify protein orientation when adsorbed to citrate AuNPs. The model generates a series of x, y, z coordinates on the GB3 surface, each with a score that predicts the localized binding propensity (**Figure 1C**, where red spheres indicate a higher score).^25^ Simplifying this model, we can reduce each GB3 variant to the total number of surface points with a score above 40 (**Table S4, Supporting Information**). No correlation was observed between this total number and either the enthalpy or free energy of binding (**Figure S4, Supporting Information**). This result suggests that, unlike for AuNPs, the folded protein structure cannot be used to predict protein binding to polystyrene surfaces.

Given that the folded structure fails to predict thermodynamic binding parameters, we sought to explore structural perturbations experimentally. To understand how the secondary structure is perturbed when bound to PSNPs, protein variants were characterized using circular dichroism (CD) spectroscopy. We focused on NF-PSNPs, because thermodynamic profiles indicated that both NF-PSNPs and COOH-PSNPs exhibited similar binding behavior. The degree of conformational change was measured by integrating the area between the curves of CD spectra in the presence and absence of NPs (**Figure 4A-C**). Traditional estimates of secondary structure fractions (e.g., using singular value decomposition) are not valid when ill-defined mixtures of ordered/disordered proteins are present,^26,48^ and we chose the integration approach because it allows for quantitative comparisons of structural perturbations across different variants. By integrating the region between 200 and 240 nm, we cover the region that is most sensitive to secondary structure changes, assessing the extent of conformational changes induced by NP binding.

**Figure 4.**
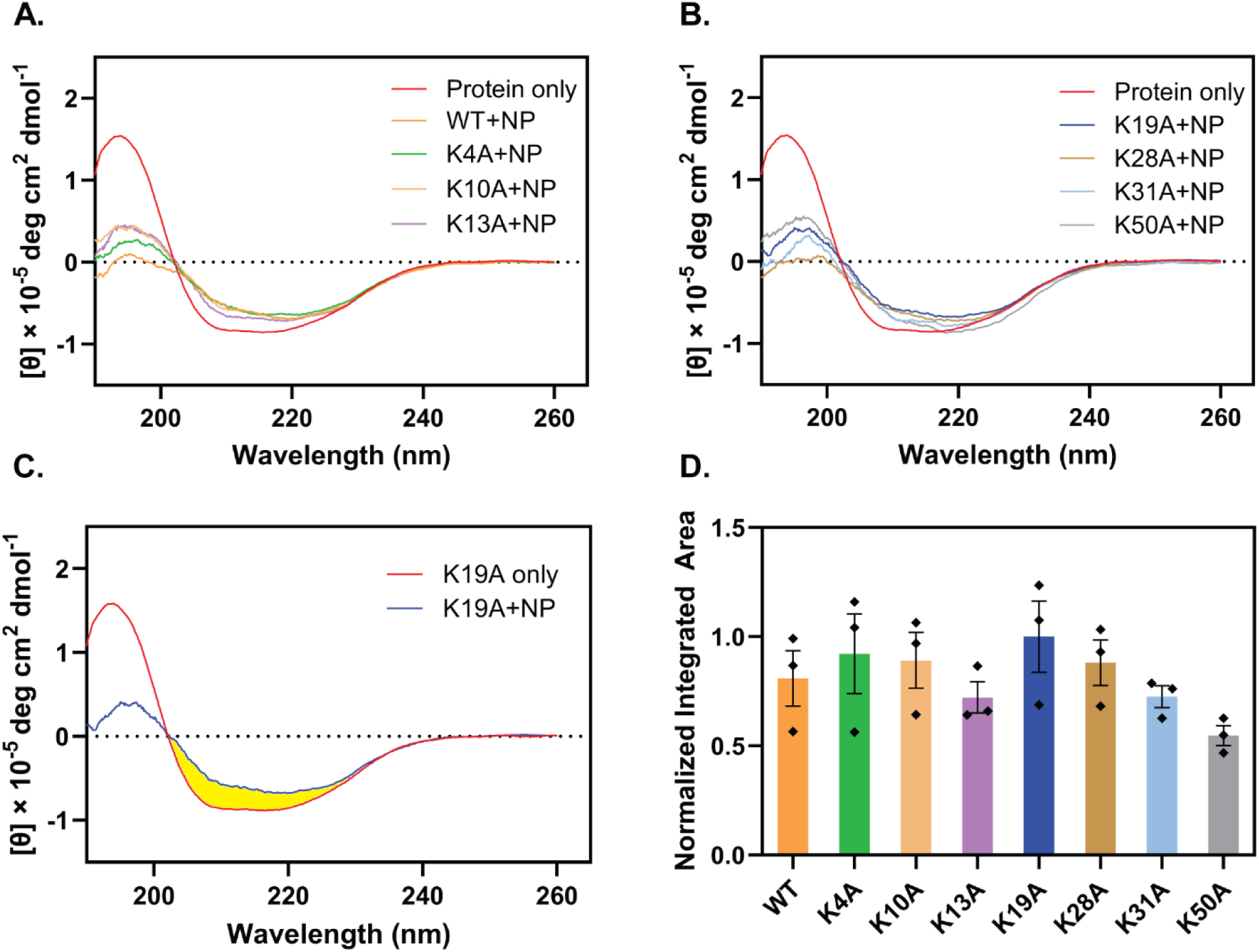
Structural perturbations of GB3 variants from CD experiments. (A, B) CD spectra of GB3 variants (A: WT, K4A, K10A and K13A; B: K19A, K28A, K31A and K50A) in the presence and absence of NF-PSNPs (C) A characteristic CD spectra of K19A in the presence and absence of PSNPs, highlighting the integrated area between the curves in the region from 200-240 nm. (D) The integrated area between the CD curves (without and without PSNPs) for GB3 variants, plotted relative to the maximum integral (K19A). Larger integrals represent a bigger difference in the CD spectrum with vs. without NF-PSNPs. Error bars represent the SEM from three sets of independently prepared samples (with/without NF-PSNPs).

The differences in CD spectra indicate that proteins undergo conformational changes upon binding to NPs. Although the K19A variant exhibited the highest structural perturbation, there was no statistically significant difference between the variants as determined using a one- way ANOVA analysis (**Figure 4D**; F(7,16) = 0.40, p = 0.89). All the variants exhibited a statistically similar degree of structural perturbation, and there is no apparent trend between any of the variants on NF-PSNPs. Additionally, no significant correlation was observed when comparing the integrated difference with the binding free energy (R^2^ = 0.076) or the binding enthalpy (R^2^ = 0.200). Thus, while ITC detects a statistically meaningful difference between the thermodynamic parameters of lysine-to-alanine variants, CD does not, although CD does indicate that structural changes are occurring.

### Protein Stability in the Presence of PSNPs is Correlated with PSNP Binding Affinity

The K19A data from ITC suggest that strongly binding proteins exhibit lower thermodynamic stability, which could lead to unfolding. To test this idea, we performed fluorescence denaturation experiments by selectively exciting the tryptophan residues at 295 nm and recording the emission spectra in the wavelength range of 305 nm to 400 nm in the presence of GdmCl as a denaturant (**Figure 5A**). Denaturation experiments were carried out on proteins in the presence and absence of NF-PSNPs and COOH-PSNPS, and the free energy of unfolding (Δ_unfold_G°) was determined using the linear extrapolation model (**Table S5, Supporting Information**)^33^. A fixed concentration of 1.5 nM PSNPs was used for all experiments, allowing relative changes in stability to be compared. The difference in free energy of unfolding (ΔΔ_unfold_G°) relative to the solution with no nanoparticles was calculated for each variant (**Figure 5B**). The ΔΔ_unfold_G° values reflect how much the polystyrene surface destabilizes the folded protein structure; large values of ΔΔ_unfold_G° should correlate with significant structural perturbations. As shown in **Figure 5C-D**, the free energy from ITC (Δ_bind_G°) and the ΔΔ_unfold_G° from folding stability experiments exhibit a positive correlation. Such a correlation reveals that the proteins with higher binding affinity are destabilized more on NPs. This phenomenon can be attributed to the competing forces involved in protein stability. While strong surface interactions may enhance binding affinity, they can also disrupt the balance of intramolecular forces that maintain the protein’s native structure. Given the complexities of measuring both Δ_bind_G° and ΔΔ_unfold_G°, it is striking that the two values correlate. Although the uncertainties are substantial, the rank order among variants is largely preserved on each type of nanoparticle. For example, K19A shows the largest Δ_bind_G°, while WT GB3, with the second largest Δ_bind_G°, displays the greatest perturbation in ΔΔ_unfold_G° for both nanoparticle types. In contrast, K10A exhibits both the lowest Δ_bind_G° and the smallest ΔΔ_unfold_G° on COOH-PSNPs, whereas on NF-PSNPs, these lowest values are observed for K31A.

**Figure 5.**
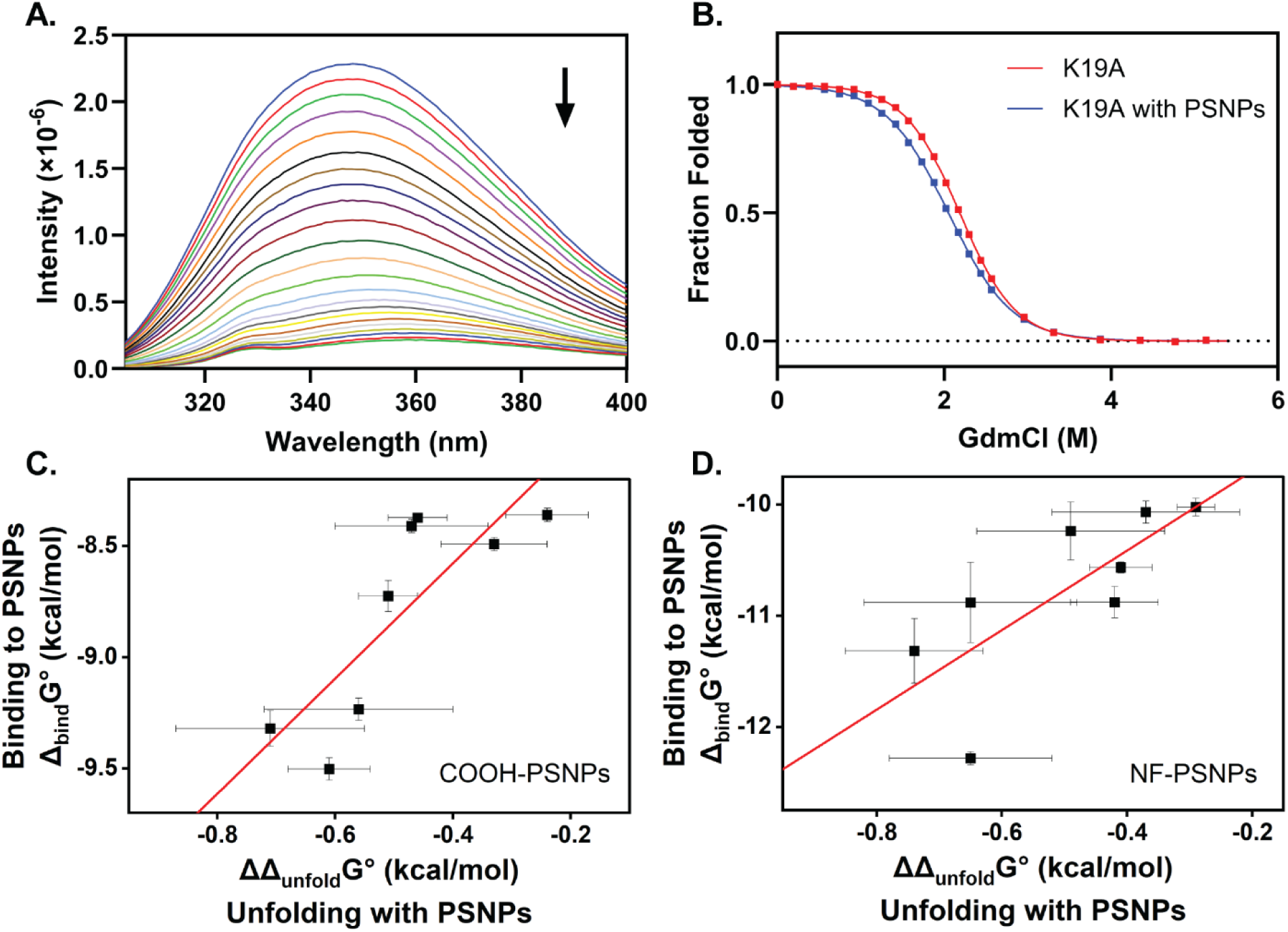
Protein Stability from fluorescence denaturation experiments. (A) A representative tryptophan fluorescence spectrum during the denaturation of GB3. The variant K19A is shown in the absence(presence/absence) of NF-PSNPs. (B) Representative normalized unfolding curves for the K19A variant in the presence and absence of NF-PSNPs. (C, D) The correlation of PSNPs binding affinity and altered protein stability in the presence of (C) COOF-PSNPs (r = 0.82, p < 0.01) (D) NF-PSNPs (r = 0.76, p < 0.03). Error bars represent SEM from three independently prepared samples (with and without PSNPs).

## Discussion

In this study, we investigate the relationship between proteins and nanoparticles using a systematic mutagenesis approach. We sought to understand the binding dynamics of GB3 variants to PSNPs, specifically addressing the influence of lysine residues on these interactions. Binding affinities and thermodynamic profiles from ITC provide a detailed view of these interactions, revealing that even subtle changes in protein structure can markedly influence adsorption.

The significant disparities in binding affinities among variants on both COOH and NF-PSNPs demonstrated this sensitivity. Particularly notable is the much higher binding affinity of K19A, which suggests a distinct mode of interaction compared to other variants. This difference is further supported by its dissimilar enthalpy change, showing a consistently exothermic profile in ITC thermograms. This finding contrasts with the exothermic-to-endothermic shift observed for other variants, indicating a unique adsorption mechanism for K19A. One might expect that removing a basic residue would reduce binding to negatively charged surfaces because of electrostatic repulsion; however, K19A binds substantially more strongly to both NF- and COOH-PSNPs.

The stronger interaction of K19A can be attributed to the location of K19 in GB3’s structure. K19 lies near the edge of a β-sheet with no nearby acidic amino acids. Mutating this residue to alanine appears to flatten the surrounding protein surface, allowing GB3 to adopt a more favorable initial orientation on the polystyrene surface, potentially unfolding after the initial binding occurs. Unlike K19, the other lysine residues are paired with nearby glutamate or aspartate residues, which likely reduces the possibility of other variants forming stable contacts with the surface. Presumably, localized electrostatic repulsions reduce the binding affinities beyond a simple loss of a positive charge. Steric factors may also play a role. This substantial experimental dataset should prove invaluable for researchers trying to validate and extend computational simulations of protein-surface interactions. For example, Sahihi *et al.* modeled the adsorption of SARS-CoV-2 spike protein on polystyrene surfaces using molecular dynamics simulations,^49^ and Hollóczki *et al*. used simulations to study protein secondary structure changes in the presence of microplastics.^50^ While informative, neither study incorporated thermodynamic parameters of the type presented here.

Additionally, the different binding affinities of the COOH- and NF-PSNPs further suggest that nanoparticle surface chemistry significantly influences binding. While this observation is not unexpected, the fact that KXA variants exhibit a different rank order on the two surfaces highlights the complexity of these interactions. Both electrostatic and hydrophobic interactions likely play a synergistic role, even if the more negatively-charged COOH-PSNPs display lower binding affinities overall. Recent work by *Desantis et al.* supports this view by showing how the spatial arrangement of charged residues modulates the binding affinities and thermal stabilities of proteins.^51^

Given that the behavior of variants does not correlate between COOH- and NF-PSNPs, the correlation between Δ_bind_G° and ΔΔ_unfold_G° on each surface is striking. This correlation suggests that protein binding is coupled to partial protein unfolding, exposing hydrophobic regions that can strengthen adsorption. The K19A variant is especially instructive: It has the strongest binding (Δ_bind_G°) and one of the largest unfolding perturbations (ΔΔ_unfold_G°). We hypothesize that such tight binding may inhibit multilayer adsorption by reducing protein-protein interactions on the nanoparticle surface. In contrast, variants with weaker binding and less unfolding may allow looser contacts, facilitating multilayer binding. These conclusions align with the recent work of Somarathne *et al.*, who showed that protein dynamics and unfolding allow proteins to form disordered nanoparticle coronas. Hydrogen-deuterium exchange and mass spectrometry data confirm that GB3 can unfold substantially on polystyrene surfaces.^52^

Finally, the correlation observed between protein stability and PSNP binding represents a step towards predicting protein-surface binding energetics. If this relationship holds more generally, it may be possible to estimate the relative rank-order of binding using fluorescence denaturation, a less expensive method than ITC. Fluorescence denaturation experiments also avoid the high nanoparticle concentrations that often lead to aggregation in calorimetric experiments. Fluorescence studies could also be extended to larger particles, such as microplastics, to improve our understanding of protein interactions with these pollutants.^53,54^ Future experiments integrating folding stability, binding energetics, and heat capacity measurements^55,56^ may help quantify the extent of protein unfolding on polystyrene surfaces. Together, such studies could reveal the fundamental structural determinants of how proteins interact with surfaces in complex biological settings.

## Conclusions

In this study, we demonstrated that lysine-to-alanine variants of GB3 exhibit distinct binding behaviors upon interaction with carboxylate and non-functionalized PSNPs, with the K19A variant showing particularly unique characteristics. This is the first experiment of its kind, where a systematic mutagenesis study was used to explore the thermodynamics of nanoparticle binding and protein unfolding. A comprehensive analysis using ITC, NMR, CD spectroscopy, and fluorescence denaturation experiments reveals differences between variants with respect to structure, multilayer binding, and the strength of enthalpic interactions. The notable difference between K19A and other variants in their binding patterns suggests a fundamental mechanistic distinction, where K19A forms a stable monolayer while other variants participate in multilayer corona formation.

Our findings also reveal an important correlation between protein binding affinity and structural stability on polystyrene surfaces, where binding appears to be thermodynamically controlled. The observation that stronger-binding variants experience greater destabilization demonstrates that strong protein-surface interactions can lead to a correspondingly strong destabilization of structure. This is in contrast to kinetically controlled surface binding (e.g., AuNPs), where protein adsorption may be sufficiently fast to preserve the folded structure^25^. This correlation has implications for the design of nanoparticle-based systems in biomedical applications, where maintaining protein stability is crucial for functionality. The correlation between binding and protein stability may also be a useful tool for estimating surface binding energies when ITC experiments are impractical, such as in studies of larger microplastic particles.

Finally, these thermodynamic and structural data provide a useful experimental resource for validating computational simulations of protein-surface and protein-nanoparticle interactions. The experimental platform developed here enables a systematic approach for studying the complex mixture of intermolecular forces involved in protein adsorption. Future work will extend these findings to other proteins, and understanding these fundamental principles will be crucial for developing predictive models of protein-surface interactions, including the nanoparticle corona, nano- and microplastics, and fouling of macroscopic surfaces.

## Supporting Information

The Supporting Information is available free of charge at …

## Author Information

### Author Contributions

C.S.K. and N.C.F. conceived the idea and designed the experiments. C.S.K., R.P.S., N.C.H., and N.C.F. developed the methodology. C.S.K., N.C.H., and N.C.F performed the experiments. C.S.K. and N.C.F. analyzed the results. The manuscript was written through the contributions of all authors. All authors have given approval to the final version of the manuscript.

### Notes

The authors declare no competing financial interests.

## Acknowledgments

We thank Railey S. Mayatt and Rebecca A. Conner for helping with the preparation of buffers for experiments. This work was supported by the National Science Foundation under awards OIA 2414443, and CBET 2405018. Support for the MSU NMR Facility was provided by the National Science Foundation under awards CHE/MCB 2304919 and DBI 2215258.

## References

(1) Lundqvist, M.; Stigler, J.; Cedervall, T.; Berggård, T.; Flanagan, M. B.; Lynch, I.; Elia, G.; Dawson, K. The Evolution of the Protein Corona around Nanoparticles: A Test Study. ACS Nano 2011, 5 (9), 7503–7509. 10.1021/nn202458g.

(2) Mahmoudi, M.; Lynch, I.; Ejtehadi, M. R.; Monopoli, M. P.; Bombelli, F. B.; Laurent, S. Protein−Nanoparticle Interactions: Opportunities and Challenges. Chem. Rev. 2011, 111 (9), 5610–5637. 10.1021/cr100440g.

(3) Chakraborty, D.; Ethiraj, K. R.; Mukherjee, A. Understanding the Relevance of Protein Corona in Nanoparticle-Based Therapeutics and Diagnostics. RSC Adv. 2020, 10 (45), 27161–27172. 10.1039/D0RA05241H.

(4) Lynch, I.; Cedervall, T.; Lundqvist, M.; Cabaleiro-Lago, C.; Linse, S.; Dawson, K. A. The Nanoparticle–Protein Complex as a Biological Entity; a Complex Fluids and Surface Science Challenge for the 21st Century. Adv. Colloid Interface Sci. 2007, 134–135, 167–174. 10.1016/j.cis.2007.04.021.

(5) Lundqvist, M.; Stigler, J.; Elia, G.; Lynch, I.; Cedervall, T.; Dawson, K. A. Nanoparticle Size and Surface Properties Determine the Protein Corona with Possible Implications for Biological Impacts. Proc. Natl. Acad. Sci. 2008, 105 (38), 14265–14270. 10.1073/pnas.0805135105.

(6) Monopoli, M. P.; Åberg, C.; Salvati, A.; Dawson, K. A. Biomolecular Coronas Provide the Biological Identity of Nanosized Materials. Nat. Nanotechnol. 2012, 7 (12), 779–786. 10.1038/nnano.2012.207.

(7) Mahon, E.; Salvati, A.; Baldelli Bombelli, F.; Lynch, I.; Dawson, K. A. Designing the Nanoparticle–Biomolecule Interface for “Targeting and Therapeutic Delivery.” J. Controlled Release 2012, 161 (2), 164–174. 10.1016/j.jconrel.2012.04.009.

(8) Milani, S.; Baldelli Bombelli, F.; Pitek, A. S.; Dawson, K. A.; Rädler, J. Reversible *versus* Irreversible Binding of Transferrin to Polystyrene Nanoparticles: Soft and Hard Corona. ACS Nano 2012, 6 (3), 2532–2541. 10.1021/nn204951s.

(9) Vroman, L.; Adams, A.; Fischer, G.; Munoz, P. Interaction of High Molecular Weight Kininogen, Factor XII, and Fibrinogen in Plasma at Interfaces. Blood 1980, 55 (1), 156–159. 10.1182/blood.V55.1.156.156.

(10) Vilaseca, P.; Dawson, K. A.; Franzese, G. Understanding and Modulating the Competitive Surface-Adsorption of Proteins through Coarse-Grained Molecular Dynamics Simulations. Soft Matter 2013, 9 (29), 6978. 10.1039/c3sm50220a.

(11) Ducoli, S.; Federici, S.; Nicsanu, R.; Zendrini, A.; Marchesi, C.; Paolini, L.; Radeghieri, A.; Bergese, P.; Depero, L. E. A Different Protein Corona Cloaks “True-to-Life” Nanoplastics with Respect to Synthetic Polystyrene Nanobeads. Environ. Sci. Nano 2022, 9 (4), 1414–1426. 10.1039/D1EN01016F.

(12) Walczyk, D.; Bombelli, F. B.; Monopoli, M. P.; Lynch, I.; Dawson, K. A. What the Cell “Sees” in Bionanoscience. J. Am. Chem. Soc. 2010, 132 (16), 5761–5768. 10.1021/ja910675v.

(13) Fleischer, C. C.; Payne, C. K. Nanoparticle–Cell Interactions: Molecular Structure of the Protein Corona and Cellular Outcomes. Acc. Chem. Res. 2014, 47 (8), 2651–2659. 10.1021/ar500190q.

(14) Park, J.; Park, S. J.; Park, J. Y.; Kim, S.; Kwon, S.; Jung, Y.; Khang, D. Unfolded Protein Corona Surrounding Nanotubes Influence the Innate and Adaptive Immune System. Adv. Sci. 2021, 8 (8), 2004979. 10.1002/advs.202004979.

(15) Tran, T. T.; Roffler, S. R. Interactions between Nanoparticle Corona Proteins and the Immune System. Curr. Opin. Biotechnol. 2023, 84, 103010. 10.1016/j.copbio.2023.103010.

(16) Cedervall, T.; Lynch, I.; Lindman, S.; Berggård, T.; Thulin, E.; Nilsson, H.; Dawson, K. A.; Linse, S. Understanding the Nanoparticle–Protein Corona Using Methods to Quantify Exchange Rates and Affinities of Proteins for Nanoparticles. Proc. Natl. Acad. Sci. 2007, 104 (7), 2050–2055. 10.1073/pnas.0608582104.

(17) Kelly, P. M.; Åberg, C.; Polo, E.; O’Connell, A.; Cookman, J.; Fallon, J.; Krpetić, Ž.; Dawson, K. A. Mapping Protein Binding Sites on the Biomolecular Corona of Nanoparticles. Nat. Nanotechnol. 2015, 10 (5), 472–479. 10.1038/nnano.2015.47.

(18) Köditz, J.; Ulbrich-Hofmann, R.; Arnold, U. Probing the Unfolding Region of Ribonuclease A by Site-Directed Mutagenesis. Eur. J. Biochem. 2004, 271 (20), 4147–4156. 10.1111/j.1432-1033.2004.04355.x.

(19) Hughson, F. M.; Barrick, D.; Baldwin, R. L. Probing the Stability of a Partly Folded Apomyoglobin Intermediate by Site-Directed Mutagenesis. Biochemistry 1991, 30 (17), 4113–4118. 10.1021/bi00231a001.

(20) Shoichet, B. K.; Baase, W. A.; Kuroki, R.; Matthews, B. W. A Relationship between Protein Stability and Protein Function. Proc. Natl. Acad. Sci. 1995, 92 (2), 452–456. 10.1073/pnas.92.2.452.

(21) Derrick, J. P.; Wigley, D. B. The Third IgG-Binding Domain from Streptococcal Protein G: An Analysis by X-Ray Crystallography of the Structure Alone and in a Complex with Fab. J. Mol. Biol. 1994, 243 (5), 906–918. 10.1006/jmbi.1994.1691.

(22) Wang, A.; Vangala, K.; Vo, T.; Zhang, D.; Fitzkee, N. C. A Three-Step Model for Protein–Gold Nanoparticle Adsorption. J. Phys. Chem. C 2014, 118 (15), 8134–8142. 10.1021/jp411543y.

(23) Wang, A.; Perera, Y. R.; Davidson, M. B.; Fitzkee, N. C. Electrostatic Interactions and Protein Competition Reveal a Dynamic Surface in Gold Nanoparticle–Protein Adsorption. J. Phys. Chem. C 2016, 120 (42), 24231–24239. 10.1021/acs.jpcc.6b08469.

(24) Xu, J. X.; Alom, Md. S.; Fitzkee, N. C. Quantitative Measurement of Multiprotein Nanoparticle Interactions Using NMR Spectroscopy. Anal. Chem. 2021, 93 (35), 11982–11990. 10.1021/acs.analchem.1c01911.

(25) Xu, J. X.; Alom, Md. S.; Yadav, R.; Fitzkee, N. C. Predicting Protein Function and Orientation on a Gold Nanoparticle Surface Using a Residue-Based Affinity Scale. Nat. Commun. 2022, 13 (1), 7313. 10.1038/s41467-022-34749-w.

(26) Somarathne, R. P.; Amarasekara, D. L.; Kariyawasam, C. S.; Robertson, H. A.; Mayatt, R.; Gwaltney, S. R.; Fitzkee, N. C. Protein Binding Leads to Reduced Stability and Solvated Disorder in the Polystyrene Nanoparticle Corona. Small 2024, 2305684. 10.1002/smll.202305684.

(27) Somarathne, R. P.; Chappell, E. R.; Perera, Y. R.; Yadav, R.; Park, J. Y.; Fitzkee, N. C. Understanding How Staphylococcal Autolysin Domains Interact With Polystyrene Surfaces. Front. Microbiol. 2021, 12, 658373. 10.3389/fmicb.2021.658373.

(28) Winzen, S.; Schwabacher, J. C.; Müller, J.; Landfester, K.; Mohr, K. Small Surfactant Concentration Differences Influence Adsorption of Human Serum Albumin on Polystyrene Nanoparticles. Biomacromolecules 2016, 17 (11), 3845–3851. 10.1021/acs.biomac.6b01503.

(29) Berry, J. D.; Neeson, M. J.; Dagastine, R. R.; Chan, D. Y. C.; Tabor, R. F. Measurement of Surface and Interfacial Tension Using Pendant Drop Tensiometry. J. Colloid Interface Sci. 2015, 454, 226–237. 10.1016/j.jcis.2015.05.012.

(30) Scheuermann, T. H.; Brautigam, C. A. High-Precision, Automated Integration of Multiple Isothermal Titration Calorimetric Thermograms: New Features of NITPIC. Methods 2015, 76, 87–98. 10.1016/j.ymeth.2014.11.024.

(31) Le, V. H.; Buscaglia, R.; Chaires, J. B.; Lewis, E. A. Modeling Complex Equilibria in Isothermal Titration Calorimetry Experiments: Thermodynamic Parameters Estimation for a Three-Binding-Site Model. Anal. Biochem. 2013, 434 (2), 233–241. 10.1016/j.ab.2012.11.030.

(32) Grimsley, G. R.; Huyghues-Despointes, B. M. P.; Pace, C. N.; Scholtz, J. M. Preparation of Urea and Guanidinium Chloride Stock Solutions for Measuring Denaturant-Induced Unfolding Curves. Cold Spring Harb. Protoc. 2006, 2006 (1), pdb.prot4241. 10.1101/pdb.prot4241.

(33) Pace, C. N.; Shaw, K. L. Linear Extrapolation Method of Analyzing Solvent Denaturation Curves. Proteins Struct. Funct. Genet. 2000, 41 (S4), 1–7. 10.1002/1097-0134(2000)41:4+<1::AID-PROT10>3.0.CO;2-2.

(34) Lakomek, N.-A.; Ying, J.; Bax, A. Measurement of 15N Relaxation Rates in Perdeuterated Proteins by TROSY-Based Methods. J. Biomol. NMR 2012, 53 (3), 209–221. 10.1007/s10858-012-9626-5.

(35) Ceccon, A.; Tugarinov, V.; Clore, G. M. TiO_2_ Nanoparticles Catalyze Oxidation of Huntingtin Exon 1-Derived Peptides Impeding Aggregation: A Quantitative NMR Study of Binding and Kinetics. J. Am. Chem. Soc. 2019, 141 (1), 94–97. 10.1021/jacs.8b11441.

(36) Ceccon, A.; Tugarinov, V.; Boughton, A. J.; Fushman, D.; Clore, G. M. Probing the Binding Modes of a Multidomain Protein to Lipid-Based Nanoparticles by Relaxation-Based NMR. J. Phys. Chem. Lett. 2017, 8 (11), 2535–2540. 10.1021/acs.jpclett.7b01019.

(37) Kihara, S.; Van Der Heijden, N. J.; Seal, C. K.; Mata, J. P.; Whitten, A. E.; Köper, I.; McGillivray, D. J. Soft and Hard Interactions between Polystyrene Nanoplastics and Human Serum Albumin Protein Corona. Bioconjug. Chem. 2019, 30 (4), 1067–1076. 10.1021/acs.bioconjchem.9b00015.

(38) Weber, C.; Simon, J.; Mailänder, V.; Morsbach, S.; Landfester, K. Preservation of the Soft Protein Corona in Distinct Flow Allows Identification of Weakly Bound Proteins. Acta Biomater. 2018, 76, 217–224. 10.1016/j.actbio.2018.05.057.

(39) Prozeller, D.; Morsbach, S.; Landfester, K. Isothermal Titration Calorimetry as a Complementary Method for Investigating Nanoparticle-Protein Interactions. Nanoscale 2019, 11 (41). 10.1039/c9nr05790k.

(40) Baker, N. A.; Sept, D.; Joseph, S.; Holst, M. J.; McCammon, J. A. Electrostatics of Nanosystems: Application to Microtubules and the Ribosome. Proc. Natl. Acad. Sci. 2001, 98 (18), 10037– 10041. 10.1073/pnas.181342398.

(41) Viola, G.; Barracchia, C. G.; Tira, R.; Parolini, F.; Leo, G.; Bellanda, M.; Munari, F.; Capaldi, S.; D’Onofrio, M.; Assfalg, M. New Paradigm for Nano–Bio Interactions: Multimolecular Assembly of a Prototypical Disordered Protein with Ultrasmall Nanoparticles. Nano Lett. 2022, 22 (22), 8875–8882. 10.1021/acs.nanolett.2c02902.

(42) De, M.; You, C.-C.; Srivastava, S.; Rotello, V. M. Biomimetic Interactions of Proteins with Functionalized Nanoparticles: A Thermodynamic Study. J. Am. Chem. Soc. 2007, 129 (35), 10747–10753. 10.1021/ja071642q.

(43) Balog, S.; Rodriguez-Lorenzo, L.; Monnier, C. A.; Obiols-Rabasa, M.; Rothen-Rutishauser, B.; Schurtenberger, P.; Petri-Fink, A. Characterizing Nanoparticles in Complex Biological Media and Physiological Fluids with Depolarized Dynamic Light Scattering. Nanoscale 2015, 7 (14), 5991–5997. 10.1039/C4NR06538G.

(44) García-Álvarez, R.; Vallet-Regí, M. Hard and Soft Protein Corona of Nanomaterials: Analysis and Relevance. Nanomaterials 2021, 11 (4), 888. 10.3390/nano11040888.

(45) Anthis, N. J.; Clore, G. M. Visualizing Transient Dark States by NMR Spectroscopy. Q. Rev. Biophys. 2015, 48 (1), 35–116. 10.1017/S0033583514000122.

(46) Randika Perera, Y.; Hill, R. A.; Fitzkee, N. C. Protein Interactions with Nanoparticle Surfaces: Highlighting Solution NMR Techniques. Isr. J. Chem. 2019, 59 (11–12), 962–979. 10.1002/ijch.201900080.

(47) Leavitt, S.; Freire, E. Direct Measurement of Protein Binding Energetics by Isothermal Titration Calorimetry. Curr. Opin. Struct. Biol. 2001, 11 (5), 560–566. 10.1016/S0959-440X(00)00248-7.

(48) Johnson, W. C. Protein Secondary Structure and Circular Dichroism: A Practical Guide. Proteins Struct. Funct. Bioinforma. 1990, 7 (3), 205–214. 10.1002/prot.340070302.

(49) Sahihi, M.; Faraudo, J. Molecular Dynamics Simulations of Adsorption of SARS-CoV-2 Spike Protein on Polystyrene Surface. J. Chem. Inf. Model. 2022, 62 (16), 3814–3824. 10.1021/acs.jcim.2c00562.

(50) Hollóczki, O.; Gehrke, S. Nanoplastics Can Change the Secondary Structure of Proteins. Sci. Rep. 2019, 9 (1), 16013. 10.1038/s41598-019-52495-w.

(51) Desantis, F.; Miotto, M.; Di Rienzo, L.; Milanetti, E.; Ruocco, G. Spatial Organization of Hydrophobic and Charged Residues Affects Protein Thermal Stability and Binding Affinity. Sci. Rep. 2022, 12 (1), 12087. 10.1038/s41598-022-16338-5.

(52) Somarathne, R. P.; Misra, S. K.; Kariyawasam, C. S.; Kessl, J. J.; Sharp, J. S.; Fitzkee, N. C. Exploring Residue-Level Interactions between the Biofilm-Driving R2ab Protein and Polystyrene Nanoparticles. Langmuir 2024, 40 (2), 1213–1222. 10.1021/acs.langmuir.3c02609.

(53) Gligorijevic, N.; Lujic, T.; Mutic, T.; Vasovic, T.; De Guzman, M. K.; Acimovic, J.; Stanic-Vucinic, D.; Cirkovic Velickovic, T. Ovalbumin Interaction with Polystyrene and Polyethylene Terephthalate Microplastics Alters Its Structural Properties. Int. J. Biol. Macromol. 2024, 267, 131564. 10.1016/j.ijbiomac.2024.131564.

(54) Ghosal, S.; Bag, S.; Burman, M. D.; Bhowmik, S. Multispectroscopic Investigations of the Binding Interaction between Polyethylene Microplastics and Human Hemoglobin. J. Phys. Chem. Lett. 2023, 14 (46), 10328–10332. 10.1021/acs.jpclett.3c02632.

(55) Spolar, R. S.; Record, M. T. Coupling of Local Folding to Site-Specific Binding of Proteins to DNA. Science 1994, 263 (5148), 777–784. 10.1126/science.8303294.

(56) McConnell, K. D.; Fitzkee, N. C.; Emerson, J. P. Metal Ion Binding Induces Local Protein Unfolding and Destabilizes Human Carbonic Anhydrase II. Inorg. Chem. 2022, 61 (3), 1249–1253. 10.1021/acs.inorgchem.1c03271.

