## Supporting Information for "Thermodynamic Analysis of Protein-Nanoparticle Interactions Links Binding Affinity and Structural Stability"

### Supporting Tables

#### Table S1. Characterization of polystyrene nanoparticles

| | Hydrodynamic Diameter (nm) | Zeta Potential (mV) |
|---|---|---|
| COOH-PSNPs | 58.35 ± 0.19 | -68.87 ± 0.52 |
| NF-PSNPs | 60.57 ± 0.76 | -54.43 ± 0.79 |

#### Table S2. Thermodynamic parameters for GB3 variants binding to COOH-PSNPs

| COOH | N | | $\Delta_{bind}G°$ (kcal mol$^{-1}$) | | $\Delta H°$ (kcal mol$^{-1}$) | |
|---|---|---|---|---|---|---|
| | **Process 1** | **Process 2** | **Process 1** | **Process 2** | **Process 1** | **Process 2** |
| **WT** | 308 ± 8 | 107 ± 3 | -9.33 ± 0.08 | -8.01 ± 0.11 | -24.3 ± 0.7 | 20.3 ± 3.1 |
| **K4A** | 419 ± 2 | 0.79 ± 0.24 | -8.50 ± 0.03 | -7.75 ± 0.06 | -29.8 ± 0.4 | (1.1± 0.4) $\times 10^4$ |
| **K10A** | 409 ± 2 | 0.24 ± 0.02 | -8.36 ± 0.03 | -7.69 ± 0.05 | -26.5 ± 0.8 | (1.8± 0.1) $\times 10^4$ |
| **K13A** | 412 ± 12 | 7 ± 5 | -8.38 ± 0.02 | -7.60 ± 0.04 | -27.4 ± 0.4 | (5.1± 2.4) $\times 10^3$ |
| **K19A** | 325 ± 30 | 81 ± 15 | -9.51 ± 0.05 | -8.43 ± 0.35 | -22.0 ± 1.2 | 18.3 ± 2.7 |
| **K28A** | 406 ± 10 | 0.19 ± 0.05 | -8.41 ± 0.03 | -7.67 ± 0.09 | -27.5 ± 0.8 | (2.7 ± 0.5) $\times 10^4$ |
| **K31A** | 286 ± 7 | 135 ± 2 | -9.24 ± 0.05 | -7.67 ± 0.09 | -32.2 ± 0.3 | 24.5 ± 0.6 |
| **K50A** | 386 ± 18 | 60 ± 13 | -8.73 ± 0.07 | -7.83 ± 0.14 | -35.4 ± 1.3 | 134.2 ± 60.1 |

**Table S3. Thermodynamic parameters for GB3 variants binding to NF-PSNPs**

| NF | N | | $\Delta_{bind}G°$ (kcal mol$^{-1}$) | | $\Delta H°$ (kcal mol$^{-1}$) | |
|---|---|---|---|---|---|---|
| | **Process 1** | **Process 2** | **Process 1** | **Process 2** | **Process 1** | **Process 2** |
| **WT** | 451 ± 98 | 187 ± 27 | -11.32 ± 0.29 | -9.66 ± 0.32 | -17.7 ± 1.8 | 7.0 ± 1.5 |
| **K4A** | 770 ± 8 | 0.37 ± 0.25 | -10.07 ± 0.10 | -9.13 ± 0.06 | -19.0 ± 0.9 | (6.6 ± 2.5) ×$10^4$ |
| **K10A** | 596 ± 15 | 90 ± 16 | -10.24 ± 0.26 | -9.13 ± 0.24 | -20.5 ± 0.2 | 53.2 ± 16.9 |
| **K13A** | 436 ± 14 | 56 ± 7 | -10.57 ± 0.05 | -9.40 ± 0.13 | -31.2 ± 1.5 | 78.6 ± 14.1 |
| **K19A** | 442 ± 60 | 211 ± 7 | -12.29 ± 0.06 | -10.24 ± 0.14 | -19.0 ± 0.4 | 2.7 ± 0.1 |
| **K28A** | 580 ± 21 | 184 ± 16 | -10.89 ± 0.36 | -9.35 ± 0.34 | -20.4 ± 0.6 | 11.9 ± 1.2 |
| **K31A** | 664 ± 8 | 4 ± 3 | -10.03 ± 0.08 | -9.26 ± 0.10 | -27.2 ± 0.4 | (6.7 ± 2.8) ×$10^4$ |
| **K50A** | 546 ± 26 | 243 ± 40 | -10.88 ± 0.14 | -9.12 ± 0.34 | -21.4 ± 0.3 | 10.3 ± 1.1 |

**Table S4. Surface binding model scores for GB3 lysine-to-alanine variants**

For each variant, a binding model was used to calculate regions on GB3 where the surface binding was high, according to predictions for citrate-capped AuNPs, where GB3 remains folded (see main text). The number of high-scoring (> 40) points on the surface is given for each variant, as calculated by code available at https://github.com/FitzkeeLab/citrate-aunp-predict. Representative surfaces are shown in Figure S4.

| Variant | Surface Binding Points |
|---|---|
| WT | 294.00 |
| K4A | 300.00 |
| K10A | 163.00 |
| K13A | 150.00 |
| K19A | 255.00 |
| K28A | 232.00 |
| K31A | 294.00 |
| K50A | 331.00 |

**Table S5. Protein unfolding stabilities measured in the presence and absence of PSNPs**

| Sample | $\Delta_{unfold}G°$ (kcal mol$^{-1}$) | m (kcal mol$^{-1}$ M$^{-1}$) |
|---|---|---|
| WT only | 3.92 ± 0.06 | 1.70 ± 0.02 |
| + NF-PSNPs | 3.18 ± 0.15 | 1.52 ± 0.05 |
| + COOH-PSNPs | 3.21 ± 0.04 | 1.52 ± 0.04 |
| K4A only | 2.98 ± 0.06 | 1.65 ± 0.01 |
| + NF-PSNPs | 2.46 ± 0.08 | 1.52 ± 0.03 |
| + COOH-PSNPs | 2.61 ± 0.11 | 1.61 ± 0.00 |
| K10A only | 3.30 ± 0.11 | 1.61 ± 0.04 |
| + NF-PSNPs | 2.80 ± 0.18 | 1.48 ± 0.06 |
| + COOH-PSNPs | 2.89 ± 0.02 | 1.58 ± 0.04 |
| K13A only | 3.13 ± 0.07 | 1.58 ± 0.03 |
| + NF-PSNPs | 2.72 ± 0.01 | 1.50 ± 0.01 |
| + COOH-PSNPs | 2.68 ± 0.03 | 1.33 ± 0.01 |
| K19A only | 3.86 ± 0.05 | 1.77 ± 0.02 |
| + NF-PSNPs | 3.21 ± 0.10 | 1.56 ± 0.03 |
| + COOH-PSNPs | 3.25 ± 0.06 | 1.74 ± 0.03 |
| K28A only | 3.76 ± 0.11 | 1.57 ± 0.01 |
| + NF-PSNPs | 3.11 ± 0.21 | 1.46 ± 0.06 |
| + COOH-PSNPs | 2.73 ± 0.02 | 1.16 ± 0.02 |
| K31A only | 3.05 ± 0.00 | 1.81 ± 0.03 |
| + NF-PSNPs | 2.76 ± 0.05 | 1.71 ± 0.01 |
| + COOH-PSNPs | 2.49 ± 0.23 | 1.43 ± 0.09 |
| K50A only | 3.25 ± 0.01 | 1.67 ± 0.01 |
| + NF-PSNPs | 2.83 ± 0.11 | 1.60 ± 0.04 |
| + COOH-PSNPs | 2.74 ± 0.07 | 1.51 ± 0.03 |

### Supporting Figures

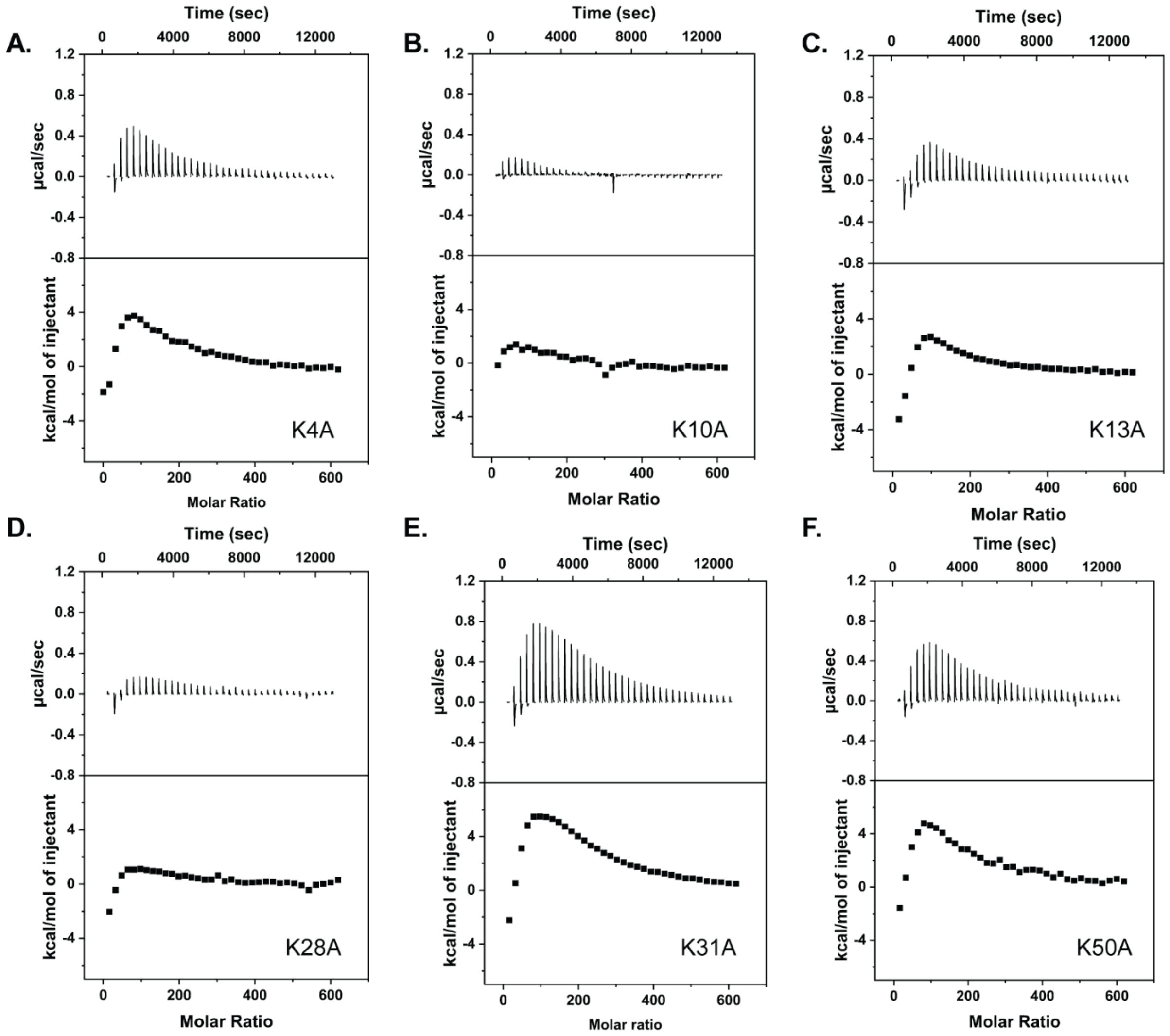

**Figure S1. Evidence for multilayer binding in non-K19A GB3 variants.**

ITC thermograms of binding after removal of weakly bound proteins for (A) K4A GB3 and (B) K10A GB3 (C) K13A (D) K28A (E) K31A (F) K50A.

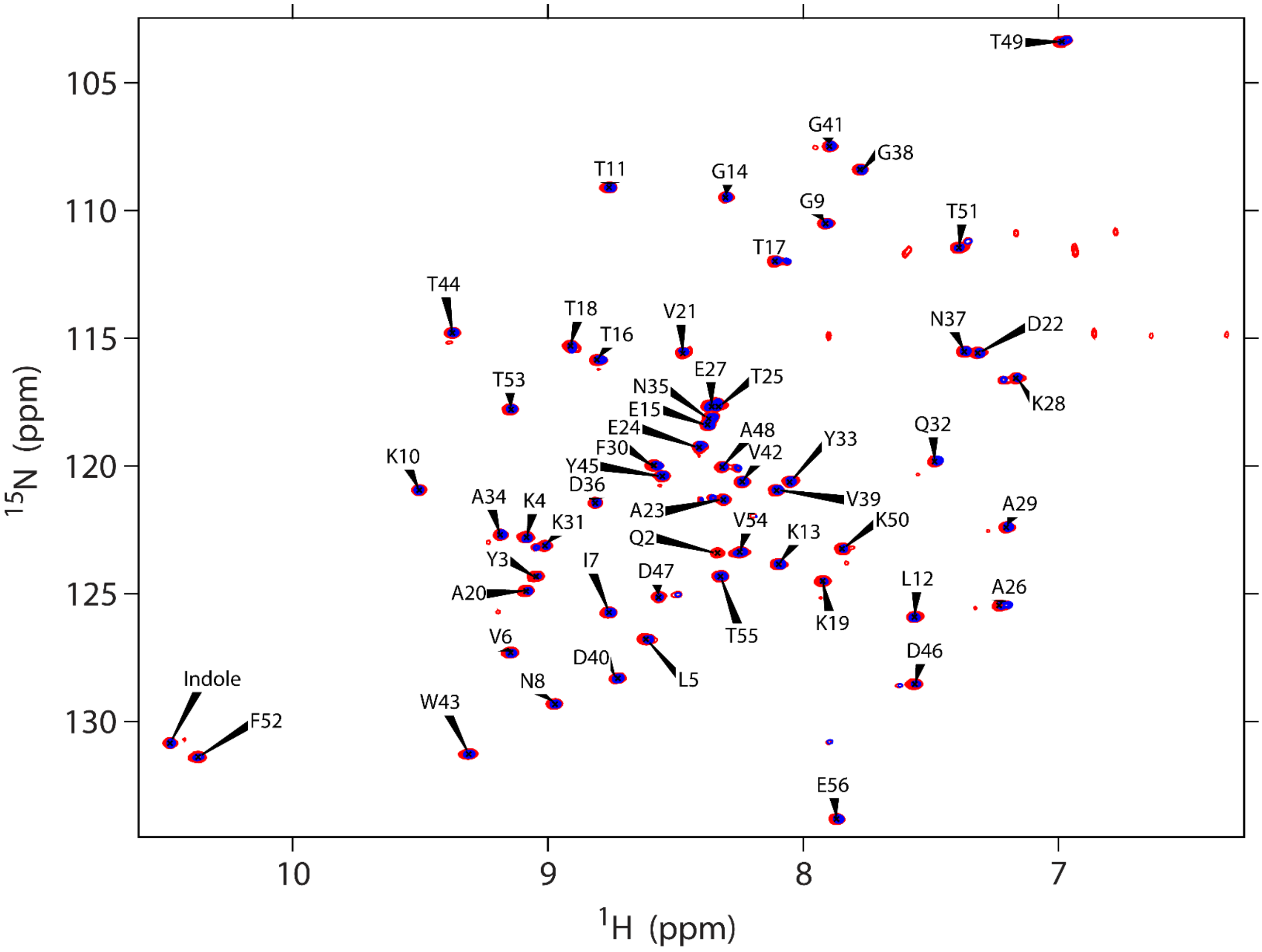

**Figure S2. Overlay of HSQC NMR spectra of WTGB3 in the absence and presence of PSNPs in T1ρ NMR experiments**

$^{1}$H-$^{15}$N HSQC NMR spectra of WTGB3 in the absence (red) and presence (blue) of PSNPs at 2 ms delay time.

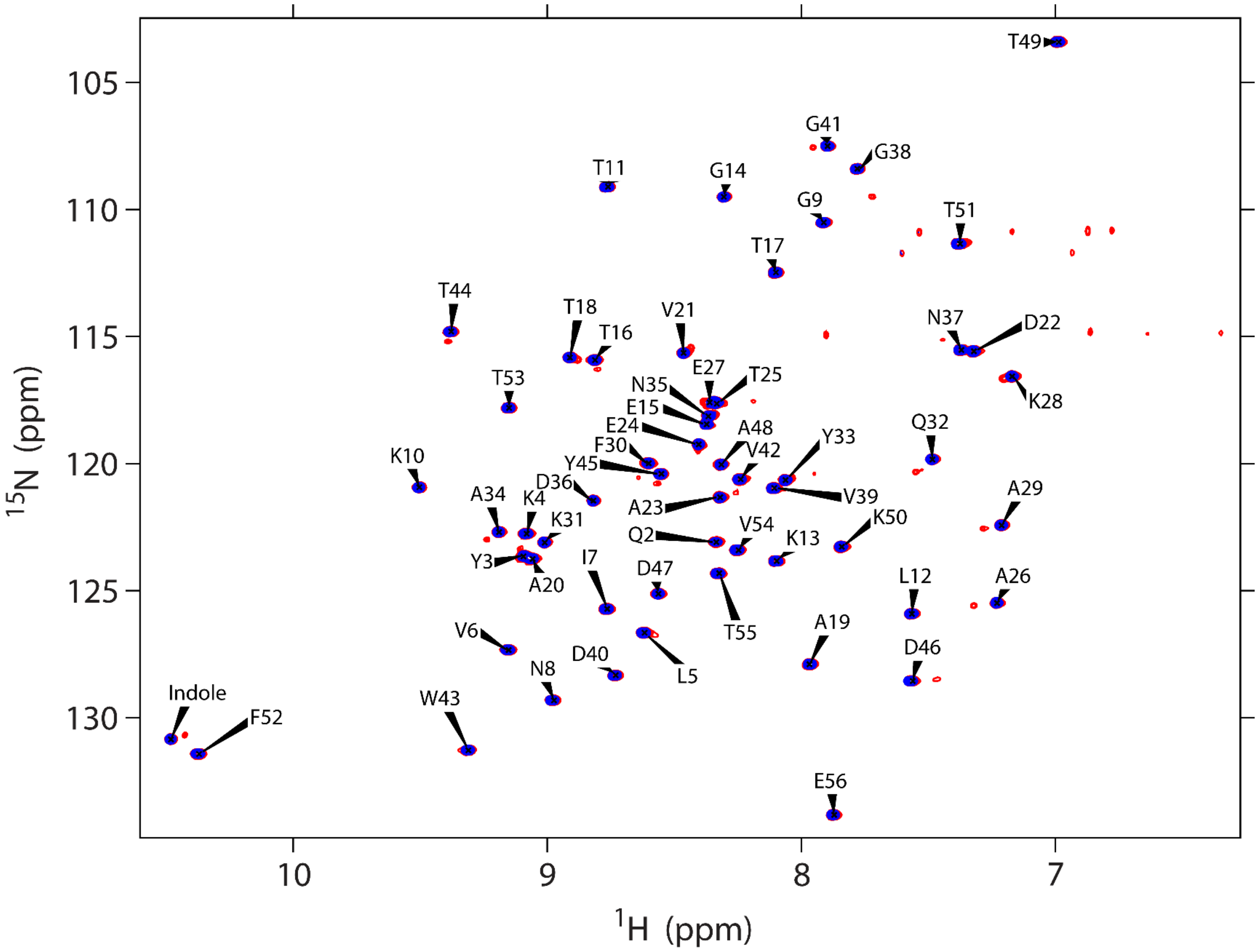

**Figure S3. Overlay of HSQC NMR spectra of K19AGB3 in the absence and presence of PSNPs in T1ρ NMR experiments**

$^{1}$H-$^{15}$N HSQC NMR spectra of K19AGB3 in the absence (red) and presence (blue) of PSNPs at 2 ms delay time.

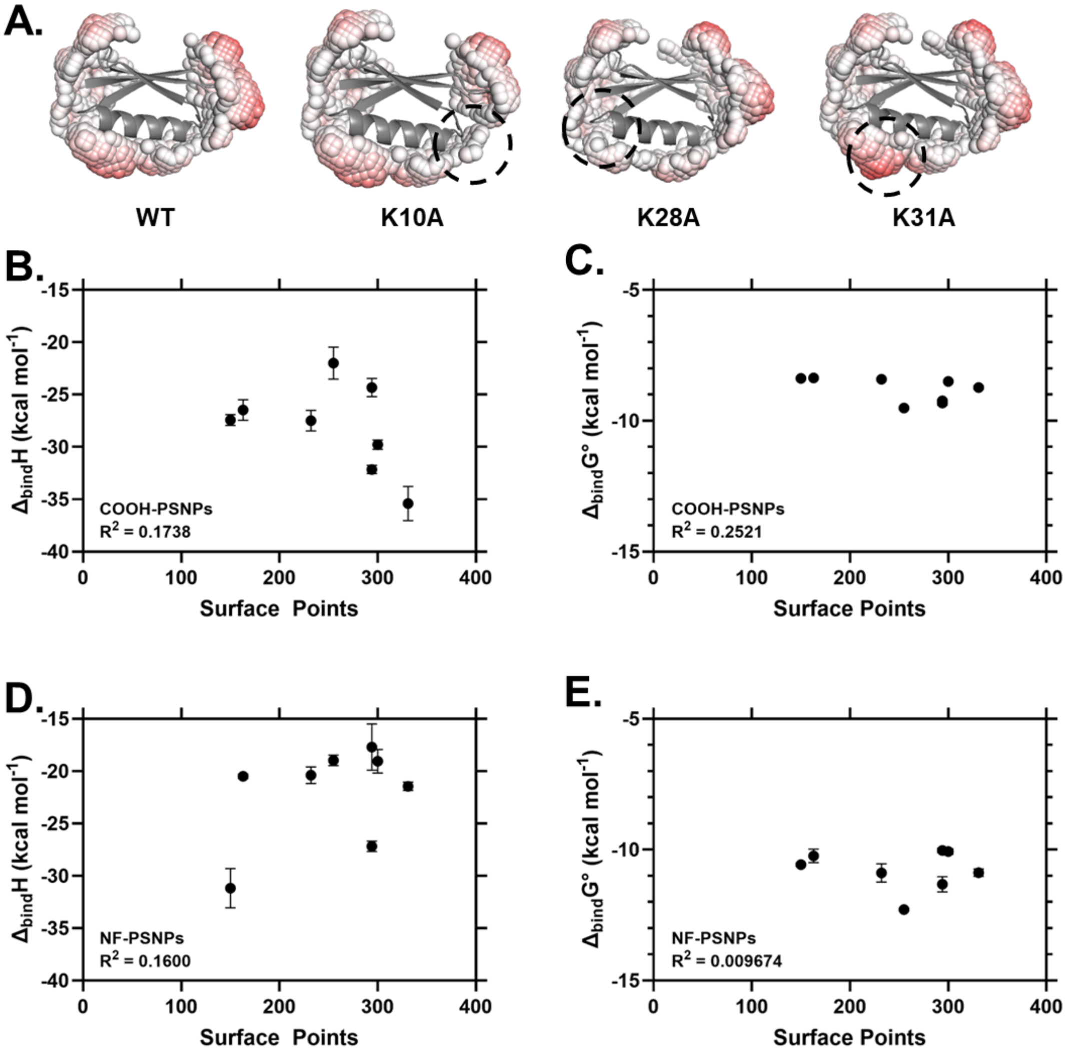

**Figure S4. Correlation between a surface binding model and calorimetric parameters**

(A.) Representative interaction surfaces predicted for GB3 lysine-to-alanine variants using a citrate-capped gold nanoparticle prediction model. Regions with high interaction potential are shown as spheres, with red spheres having the highest potential. In this model, proteins are assumed to remain folded. Differences from wild-type are annotated with a black dashed line. (B.-E.) Correlation between calorimetric parameters ($\Delta_{bind}H$ and $\Delta_{bind}G$) and surface points identified with the model shown in panel A. The number of high-potential surface points (> 40) is plotted against thermodynamic parameters for COOH-PSNPs (B., C.) and NF-PSNPs (D., E.). Correlation coefficients are poor, suggesting that these variants are not well represented by a folded, globular structure.
